# Interleukin-6 Drives Key Pathologic Outcomes in Experimental Acetaminophen-induced Liver Failure

**DOI:** 10.1101/2021.11.15.468664

**Authors:** Katherine Roth, Jenna Strickland, Romina Gonzalez-Pons, Asmita Pant, Ting-Chieh Yen, Robert Freeborn, Rebekah Kennedy, Bharat Bhushan, Allison Boss, Cheryl E. Rockwell, Anne M. Dorrance, Udayan Apte, James P. Luyendyk, Bryan L. Copple

## Abstract

**Background and Aims:** In severe cases of acetaminophen (APAP) overdose, acute liver injury rapidly progresses to acute liver failure (ALF), producing life-threatening complications including, hepatic encephalopathy (HE) and multi-organ failure (MOF). Systemic levels of interleukin-6 (IL-6) and IL-10 are highest in ALF patients with the most severe complications and the poorest prognosis. The mechanistic basis for dysregulation of these cytokines, and their association with outcome in ALF, remain poorly defined.

**Methods:** To investigate the impact of IL-6 and IL-10 in ALF, we used an experimental setting of failed liver repair after APAP overdose in which a high dose of APAP is administered (i.e., 500-600 mg/kg). Mice were treated with neutralizing antibodies to block IL-6 and IL-10.

**Results:** In mice with APAP-induced ALF, high levels of IL-10 reduced monocyte recruitment and trafficking in the liver resulting in impaired clearance of dead cell debris. Kupffer cells in these mice, displayed features of myeloid-derived suppressor cells, including high level expression of IL-10 and PD-L1, which were increased in an IL-6-dependent manner. Similar to ALF patients with HE, cerebral blood flow was reduced in mice with APAP-induced ALF. Remarkably, although IL-6 is hepatoprotective in mice treated with low doses of APAP (i.e., 300 mg/kg), IL-6 neutralization in mice with APAP-induced ALF fully restored cerebral blood flow and reduced mortality.

**Conclusion:** Collectively, these studies demonstrate that exaggerated production of IL-6 in APAP-induced ALF triggers immune suppression (i.e., high levels of IL-10 and PD-L1), reduces cerebral blood flow (a feature of hepatic encephalopathy), disrupts liver repair (i.e., failed clearance of dead cells), and increases mortality.

## Introduction

Acetaminophen (APAP) overdose remains the number one cause of acute liver failure (ALF) in the United States ^1^. In severe cases, acute liver injury rapidly progresses to acute liver failure (ALF), producing life-threatening cardiac instability, hepatic encephalopathy, and multi-organ failure ^2, 3^. First line therapy for APAP overdose is N-acetyl cysteine (NAC), which is highly efficacious when administered during the active phase of injury ^4–6^. Because of this, the efficacy of NAC quickly declines for patients that seek medical attention beyond 8 hours after APAP ingestion ^5^. For this group of patients, supportive medical care and liver transplantation are the only remaining therapeutic options ^7, 8^. Unfortunately, despite significant improvements in clinical care and emergency liver transplantation, mortality associated with ALF remains high, underscoring the importance of developing new therapeutic approaches, particularly for patients refractory to NAC therapy ^3, 8^.

ALF patients with the poorest prognosis develop systemic inflammatory response syndrome (SIRS), a condition characterized by high systemic levels of proinflammatory cytokines, including tumor necrosis factor-α (TNF-α), CC motif chemokine ligand 2 (Ccl2), and interleukin-6 (IL-6) ^9, 10^. Clinical studies have documented a clear association between high levels of the SIRS-associated cytokine, IL-6, and a worsening of hepatic encephalopathy and development of multiorgan dysfunction syndrome (MODS), suggesting a causal role for SIRS in the pathogenesis of these extrahepatic complications ^11, 12^. The severe systemic inflammation produced by SIRS often triggers activation of anti-inflammatory pathways that generate high levels of immune-suppressive cytokines (e.g., IL-10) and immunoregulatory ligands (e.g., PD-L1) ^11, 13^. This condition, referred to as compensatory anti-inflammatory response syndrome (CARS), is a state of “immune paralysis” that is associated with increased mortality in ALF patients, in part, through an increased risk of nosocomial infections. Further, it has been suggested that the severe immune suppression produced by CARS may disrupt essential pro-repair activities of immune cells, contributing to the failed liver repair that occurs in these patients ^14^. While clinical studies have demonstrated a clear association between the severity of SIRS and CARS and patient outcome in ALF, it remains to be established whether these conditions are mechanistically involved in the pathogenesis of APAP-induced ALF.

In experimental studies of APAP overdose, high systemic levels of cytokines associated with SIRS and CARS (i.e., IL-6 and IL-10) have been reported ^15–17^. In stark contrast to ALF patients, however, increased expression of these cytokines is associated with reduced liver injury and mortality. For instance, compared to wild-type mice, IL-10 knockout mice develop markedly greater liver injury that is associated with reduced survival when treated with hepatotoxic doses of APAP ranging from 150 to 300 mg/kg ^17^. Similarly, IL-6 knockout mice treated with 300 mg/kg APAP develop greater liver injury and display reduced hepatocyte proliferation when compared to wild-type mice ^15^. Although these results suggest that IL-6 and IL-10 are not mechanistically involved in the pathogenesis of ALF, they must be considered in the context of the experimental conditions. These studies were conducted using standard experimental settings of APAP hepatotoxicity in which mice were treated with doses of APAP at or near 300 mg/kg. These doses produce liver injury that is predictably repaired thereby restoring hepatic function and triggering resolution of inflammation. As a result, ALF, and its associated extrahepatic complications (e.g., MODS, hepatic encephalopathy) do not develop in these mice.

To investigate SIRS and CARS in ALF, we used a robust experimental setting of failed liver repair after APAP overdose in which a high dose of APAP is administered (i.e., 500-600 mg/kg) to wild-type mice. This strategy, originally characterized by Bhushan and colleagues, produces liver injury that fails to repair, mirroring findings reported in ALF patients with the poorest prognosis ^18^. Failed liver repair in these mice is associated with increased mortality and occurs despite plasma levels of IL-6 that are markedly higher than those in mice treated with a nonlethal dose of APAP (i.e., 300 mg/kg) ^18^. Further, myeloid cell-dependent clearance of necrotic cells from the injured liver is disrupted, suggesting a functional impairment of myeloid cell populations ^18^. Remarkably, despite the striking differences in liver repair, the severity of liver injury is not significantly different between mice treated with 300 mg/kg or 600 mg/kg APAP, mirroring the poor association between the severity of liver injury and outcome in ALF patients ^18, 19^. Collectively, the high levels of IL-6 occurring in mice treated with 600 mg/kg APAP, coupled with impaired myeloid cell activity and high mortality, suggest that SIRS and CARS may develop in these mice. To evaluate this, we used a similar dosing strategy to determine whether cytokine dysregulation, indicative of SIRS and CARS, develops in these mice and further tested the hypothesis that SIRS and CARS contribute to the pathogenesis of ALF and its associated secondary complications.

## Materials and Methods

### ANIMALS AND TREATMENTS

6-12-week-old male C57BL/6J mice and IL-10 reporter mice (IL10^tm1.1Karp^) (Jackson Laboratories) were used for all studies. Mice were housed in a 12-hour light/dark cycle under controlled temperature (18-21°C) and humidity. Food (Rodent Chow; Harlan-Teklad) and tap water were allowed *ad libitum*.

For treatment with saline, 300 mg/kg APAP (Sigma-Aldrich, St. Louis, MO), or 600 mg/kg APAP, mice were fasted for approximately 12 hours prior to injection, as described previously ^18^. In all studies, rodent chow was returned immediately after APAP challenge.

For IL-10 neutralization studies, mice were injected with 0.5 mg *InVivo*MAb anti-mouse IL-10 antibody (Bio X Cell, clone JES5-2A5) or 0.5 mg isotype control antibody (Innovative Research, Rat IgG) at 24 hours after APAP treatment. Liver and blood were collected at 72 hours after APAP treatment. For recombinant IL-10 studies, mice were injected with 5 μg recombinant mouse IL-10 (Biolegend, San Diego, CA) or sterile saline 24 hours after APAP challenge. Liver and blood were collected at 48 hours after APAP challenge. For IL-6 neutralization studies, mice were injected with 200 μg *InVivo*MAb anti-mouse IL-6 antibody (Bio X Cell, clone MP5-20F3) or *InVivo*MAb rat IgG1 isotype control (Bio X Cell, clone HRPN) at 2 hours after APAP treatment. Liver and blood were collected, and cerebral blood flow was measured 24 hours later. All studies were approved by the Michigan State University Institutional Animal Care and Use Committee.

### SAMPLE COLLECTION

Mice were anesthetized using Fatal-Plus Solution (Vortech Pharmaceuticals) or isoflurane. Blood was collected from the inferior vena cava, and the livers were removed. A portion of each liver was fixed in 10% neutral-buffered formalin. The livers were embedded in paraffin, sectioned, and stained with hematoxylin and eosin. The area of necrosis was quantified as described by us previously ^20^. Additional portions of the liver were homogenized in TRIzol Reagent (Thermo-Fisher Scientific) for RNA isolation or were snap-frozen in liquid nitrogen for subsequent sectioning and immunofluorescence staining.

### IMMUNOFLUORESCENCE

Immunofluorescence was used to detect F4/80 and CD68 as described previously ^20^. Briefly, 8μm sections were cut from frozen livers and fixed for 10 minutes in 4% formalin. The sections were then incubated in blocking buffer (10% goat serum; 1 hour) followed by incubation with either rat anti-F4/80 antibody (Bio-Rad) diluted 1:500 or rat anti-CD68 antibody (Bio-Rad) diluted 1:500 overnight at 4°C. After washing, the sections were incubated with goat anti-rat secondary antibody conjugated to Alexa Fluor 594 for 1 hour (diluted 1:500, Thermo Fisher Scientific). Proliferating cell nuclear antigen (PCNA) was detected as described previously ^20^.

### LUMINEX IMMUNOASSAY

Cytokine levels were measured in blood serum samples by using the Bio-Plex Pro assay kit (Bio-Rad) according to manufacturer’s instructions. Bead fluorescent readings were obtained using a Luminex 200 system.

### FLOW CYTOMETRY

To isolate non-parenchymal cells, mouse livers were perfused and digested with collagenase (Collagenase H, Sigma Chemical Company) as described previously ^21^. Hepatocytes were removed by centrifugation (50 *g* for 2 minutes), and non-parenchymal cells were collected from the remaining solution by centrifugation at 300 *g* for 10 minutes. Non-parenchymal cells were washed and resuspended in FACs buffer (phosphate-buffered saline, 1% fetal bovine serum). The cells were then incubated with Fc blocking buffer (BD Biosciences; diluted 1:20) for 10 minutes at 4 °C, rinsed, and then pelleted by centrifugation at 300 *g* for 5 minutes. The cells were incubated with anti-F4/80 conjugated to Alexa-488 and anti-Ly6C conjugated to phycoerythrin (PE) for 30 minutes at 4°C. All antibodies were purchased from Biolegend. For studies with IL-10 GFP reporter mice, the following antibodies were used for flow cytometry: anti-Axl (PE), anti-CD45.1 (PE/Cy7), anti-F4/80 (AF594), anti-Cx3cr1 (AF700), anti-Ly6C (APC/Cy7), anti-Cd11b (BV605), anti-Marco (APC), anti-Ccr2 (BV650), anti-PD-L1 (PerCp/Cy5.5). All antibodies were purchased from Biolegend, except anti-Marco and anti-Axl, which were purchased from Invitrogen. The fixable dye, Zombie Aqua, was used to determine cell viability. Following incubation, cells were washed twice and fixed in formalin (Sigma) for 15 minutes at 4°C. The fixed cells were washed twice and resuspended in FACs buffer. An Attune NxT flow cytometer (Life Technologies) was then used to measure fluorescence. Signal was quantified using Attune NxT software. The gating strategy for flow cytometry is shown in Supplemental Figure 1.

### REAL-TIME PCR

Total RNA was isolated from liver samples using TRIzol Reagent (Thermo-Fisher) and reverse transcribed into cDNA as described previously^22^. Real-time PCR was performed on a QuantStudio 7 Flex Real-Time PCR System (Thermo-Fisher) using the iTaq Universal SYBR green Supermix (Bio-Rad). The following primer sequences were used: Tnf-α: Forward- 5’-AGGGTCTGGGCCATAGAACT-3’, Reverse- 5’-CCACCACGCTCTTCTGTCTAC-3’; Ccl2: Forward- 5’-CCTGCTGTTCACAGTTGCC-3’, Reverse- 5’-ATTGGGATCATCTTGCTGGT3’; Il-10: Forward- 5’-TGTCAAATTCATTCATGGCCT-3’, Reverse- 5’-ATCGATTTCTCCCCTGTGAA-3’: Rpl13a: Forward- 5’-GACCTCCTCCTTTCCCAGGC-3’, Reverse- 5’-AAGTACCTGCTTGGCCACAA-3’; PD-L1: Forward- 5’-CACGCTGAAAGTCAATGCCC-3’, Reverse- 5’-AAACATCATTCGCTGTGGCG-3’; IL-6: Forward- 5’-CCCAGTGCAAGAATCCTCGT3’, Reverse- 5’-GTCATAAGGGCTCTGTGCGT-3’.

### LASER SPECKLE CONTRAST IMAGING

Mice were anesthetized with 3.5% isoflurane in oxygen. The scalp was removed and the skull was cleared of connective tissue. A thin coat of clear shiny nail polish followed by a thin coat of matte nail polish was used to reduce glare. After reducing the isoflurane to 1.5% for three minutes, the cerebral perfusion was measured over a minute using a Perimed PeriCam PSI Zoom System and PIMSoft software (Perimed Inc, Las Vegas, NV). Regions of interest (ROI) were drawn after locating the midpoint of the sagittal suture between bregma and sigma as described by Polycarpou et. al (2016). The mean perfusion of each ROI was recorded. The average between left and right frontal, parietal and occipital lobe perfusion was calculated. Total cerebral perfusion was defined as the average of frontal, parietal, occipital and pial anastomotic regions combined.

### MICROARRAY ANALYSIS

Microarray analysis on livers from APAP-treated mice was conducted as described in detail previously ^23^.

### STATISTICAL ANALYSIS

Results are presented as the mean + SEM. Data were analyzed by a one-way or two-way Analysis of Variance (ANOVA) where appropriate. Data expressed as a percentage were transformed by arcsine square root prior to analysis. Comparisons among group means were made using the Student-Newman-Keuls test. The criterion for significance was *p* < 0.05 for all studies.

## Results

### APAP-INDUCED ALF IS COUPLED TO FAILED CLEARANCE OF NECROTIC CELL DEBRIS FROM THE INJURED LIVER

In mice challenged with 300 mg/kg APAP, liver necrosis peaked approximately 24 hours after treatment (Fig. 1A and 1B). By 48 hours, extensive inflammatory infiltrates were noted within the necrotic lesions (Fig. 1C). The inflammatory cells, along with the necrotic cell debris, were largely cleared from the liver by 72 hours (Fig. 1A and 1B). In agreement with prior studies by Bhushan and colleagues ^18^, administration of 600 mg/kg APAP produced a comparable initial hepatotoxic response (Figs. 1A and 1B). At this larger dose of APAP, however, necrotic cells persisted at 72 hours, suggesting a failure of recruited monocyte-derived macrophages (MDM) to clear dead cell debris (Fig. 1B) ^24^. Consistent with this, the necrotic lesions in these mice were largely devoid of inflammatory cells, suggesting a defect in monocyte recruitment (Fig. 1D).

**Figure 1.**
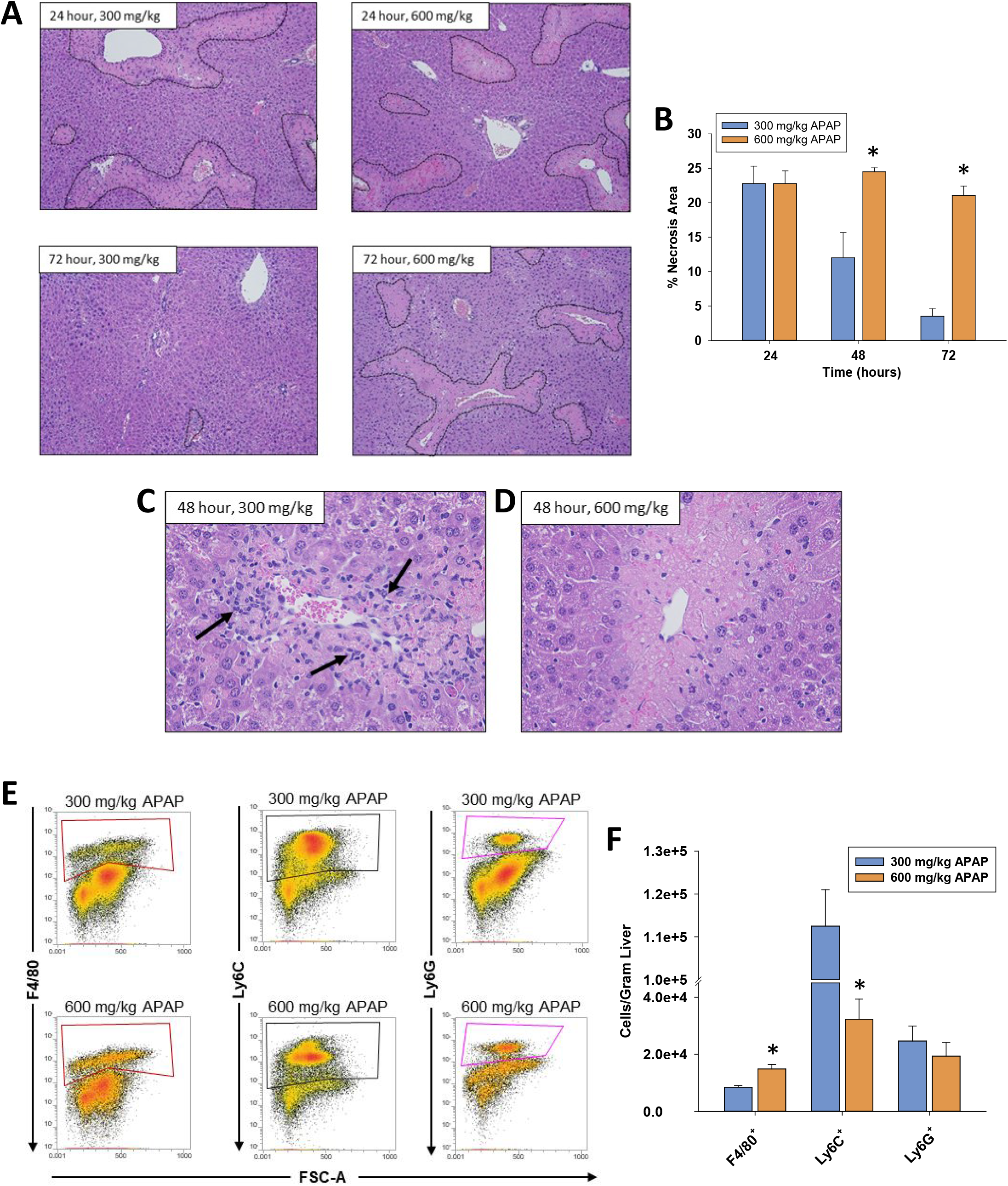
Liver injury and inflammation in APAP-treated mice. Mice were treated with either 300 mg/kg APAP or 600 mg/kg APAP for the indicated time. (A) Photomicrographs of H&E-stained liver sections. Necrotic lesions are demarcated by a dashed line. (B) Area of necrosis was quantified in sections of liver. (C,D) Photomicrographs of H&E-stained liver sections. Arrows indicate inflammatory cells within the necrotic lesions. (E) At 24 hours after APAP treatment, flow cytometry was used to detect F4/80^+^, Ly6C^+^, and Ly6G^+^ cells in the liver. Boxes indicate the positive gate. (F) Absolute cell counts from flow cytometry. *Significantly different from mice treated with 300 mg/kg APAP. All data are expressed as mean ± SEM; n = 5-10 mice per group.

### FAILED TRAFFICKING OF MDMS INTO NECROTIC LESIONS IN MICE WITH ALF

Because myeloid cells are responsible for clearing dead cell debris from the APAP-injured liver, we quantified these cells by flow cytometry ^24^. Gating for flow cytometry is shown in Supplemental Figure 1. As shown in Figures 1E and 1F, the number of Ly6G^+^ cells (i.e., neutrophils) in the liver was unaffected by APAP dose. By contrast, the number of Ly6C^+^ cells (i.e., recruited MDMs; ^25^) was substantially lower in mice treated with 600 mg/kg APAP, whereas the number of F4/80^+^ cells (i.e., Kupffer cells) was modestly higher (Fig. 1E and 1F).

Next, we quantified CD68^+^ macrophages in the liver by immunofluorescence labeling. Prior studies revealed that CD68^+^ macrophages accumulate in the livers of patients with APAP-induced ALF, and our prior studies indicate that these cells are the same as Ly6C^+^ MDMs ^26–28^. In mice treated with 300 mg/kg APAP, CD68^+^ macrophages began to accumulate in the liver and traffic into the necrotic lesions by 24 hours (Fig. 2A, necrotic lesions demarcated by white dotted lines). By 48 hours, the necrotic lesions were filled with CD68^+^ macrophages (Fig. 2A). Interestingly, fewer CD68^+^ macrophages accumulated in the livers of mice treated with 600 mg/kg APAP, and those that were present, failed to traffic into the necrotic lesions at any time point examined (Fig. 2A and 2B). We recently showed that in mice treated with 300 mg/kg APAP that macrophages within the necrotic lesions begin to express the mature tissue macrophage marker, F4/80, coincident with clearance of necrotic cells ^28^. In mice treated with 300 mg/kg APAP, F4/80^+^ macrophages were largely present outside of the necrotic lesions at 24 and 48 hours after treatment (Fig. 2C). By 72 hours, however, F4/80^+^ macrophages filled the necrotic lesions (Fig. 2C), in agreement with our prior studies ^28^. By contrast, F4/80^+^ macrophages were rarely observed in the lesions of mice treated with 600 mg/kg APAP (Fig. 2C).

**Figure 2.**
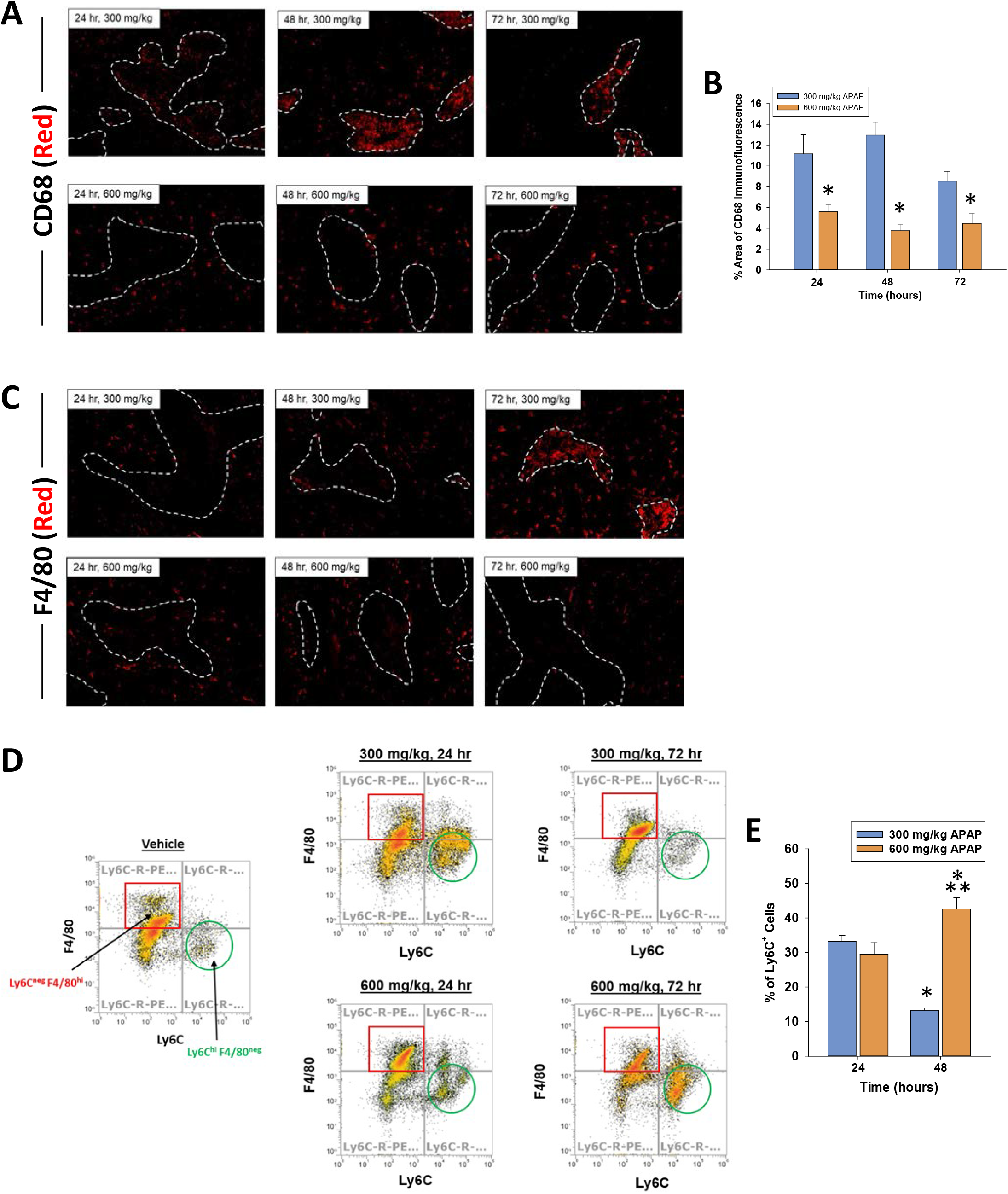
Accumulation of myeloid cells in the livers of mice treated with APAP. Mice were treated with either 300 mg/kg APAP or 600 mg/kg APAP. (A) Photomicrographs of CD68 immunofluorescent staining in liver sections. Positive staining appears red. Necrotic lesions are demarcated by a dashed white line. (B) The area of CD68 immunofluorescent staining was quantified in sections of liver. *Significantly different from mice treated with 300 mg/kg APAP. Data are expressed as mean ± SEM; n = 5 mice per group. (C) F4/80 was detected by immunofluorescence in liver sections from APAP treated mice. Treatment and time point are indicated on the photomicrograph. The necrotic lesions are demarcated by a white dashed line. F4/80^+^ macrophages appear red in the photomicrographs. Representative from an n = 5-10 mice per group. (D) Mice were treated with either vehicle control, 300 mg/kg APAP, or 600 mg/kg APAP. After 24 hours, livers were digested and Ly6C^+^ and F4/80^+^ cells were detected by flow cytometry in vehicle-treated mice. After 24 and 72 hours Ly6C^+^ and F4/80^+^ were detected in the livers of mice treated with 300 mg/kg or 600 mg/kg APAP. (E) The percentage of Ly6C^+^ cells were quantified by flow cytometry. *Significantly different from 24 hours. **Significantly different from mice treated with 300 mg/kg APAP 48 hours earlier. Data are expressed as mean ± SEM; n = 3.

Prior lineage tracing studies showed that Ly6C^+^ MDMs, which also express CD68, accumulate in the liver after APAP overdose (i.e., 300 mg/kg) and begin to express F4/80 as necrotic cells are cleared from the liver ^25^. This was associated with a shift in phenotype from proinflammatory to pro-repair typified by increased F4/80 expression and reduced Ly6C expression ^25^. As shown in Figure 2D, macrophages in the livers of vehicle-treated mice were largely F4/80^+^ Ly6C^-^ (red square). By 24 hours after treatment with 300 mg/kg APAP, however, a population of Ly6C^+^ F4/80^-^ MDMs appeared in the liver (Fig 2D, green circle). By 72 hours, this population of MDMs was reduced, indicating a transition to Ly6C^-^ F4/80^+^ macrophages (Fig. 2D) in agreement with prior studies ^25^. Ly6C^+^ F4/80^-^ MDMs also appeared in the liver 24 hours after treatment with 600 mg/kg APAP (Fig 2D, green circle). Interestingly, however, this population persisted 72 hours after treatment with 600 mg/kg APAP (Fig. 2D, green circle). Quantification revealed a similar percentage of Ly6C^+^ MDMs in the liver at 24 hours after treatment (Fig. 2E). By 72 hours, however, this population of MDMs was decreased in mice treated with 300 mg/kg APAP, whereas this population of macrophages continued to increase in mice treated with 600 mg/kg APAP (Fig. 2E). Collectively, these results suggest that Ly6C^+^ proinflammatory macrophages fail to shift phenotype in mice treated with 600 mg/kg APAP.

### SUSTAINED CYTOKINE PRODUCTION IN MICE WITH APAP-INDUCED ALF

The transition of MDMs from a proinflammatory to a pro-reparative phenotype in the APAP-injured liver is associated with a decrease in proinflammatory cytokine expression ^25, 28^. Consistent with this, in mice treated with 300 mg/kg APAP, hepatic mRNA levels of the proinflammatory cytokines, Ccl2 and Tnf-α, peaked at 24 hours and returned to baseline by 72 hours (Fig. 3A and 3B). By contrast, in mice treated with 600 mg/kg APAP, mRNA levels of these cytokines were increased by 24 hours and remained elevated at 72 hours (Fig. 3A and 3B). In addition to these cytokines, comparison of global gene expression revealed higher levels of several immunomodulatory genes in mice treated with 600 mg/kg APAP when compared to mice treated with 300 mg/kg APAP (Fig. 3C). In confirmation of the mRNA findings, serum levels of CCL2, TNF-α, interleukin-1β (IL-1β), interferon-γ (IFN-γ), and IL-4 protein were greater in mice treated with 600 mg/kg APAP at 72 hours (Fig. 3D-3H).

**Figure 3.**
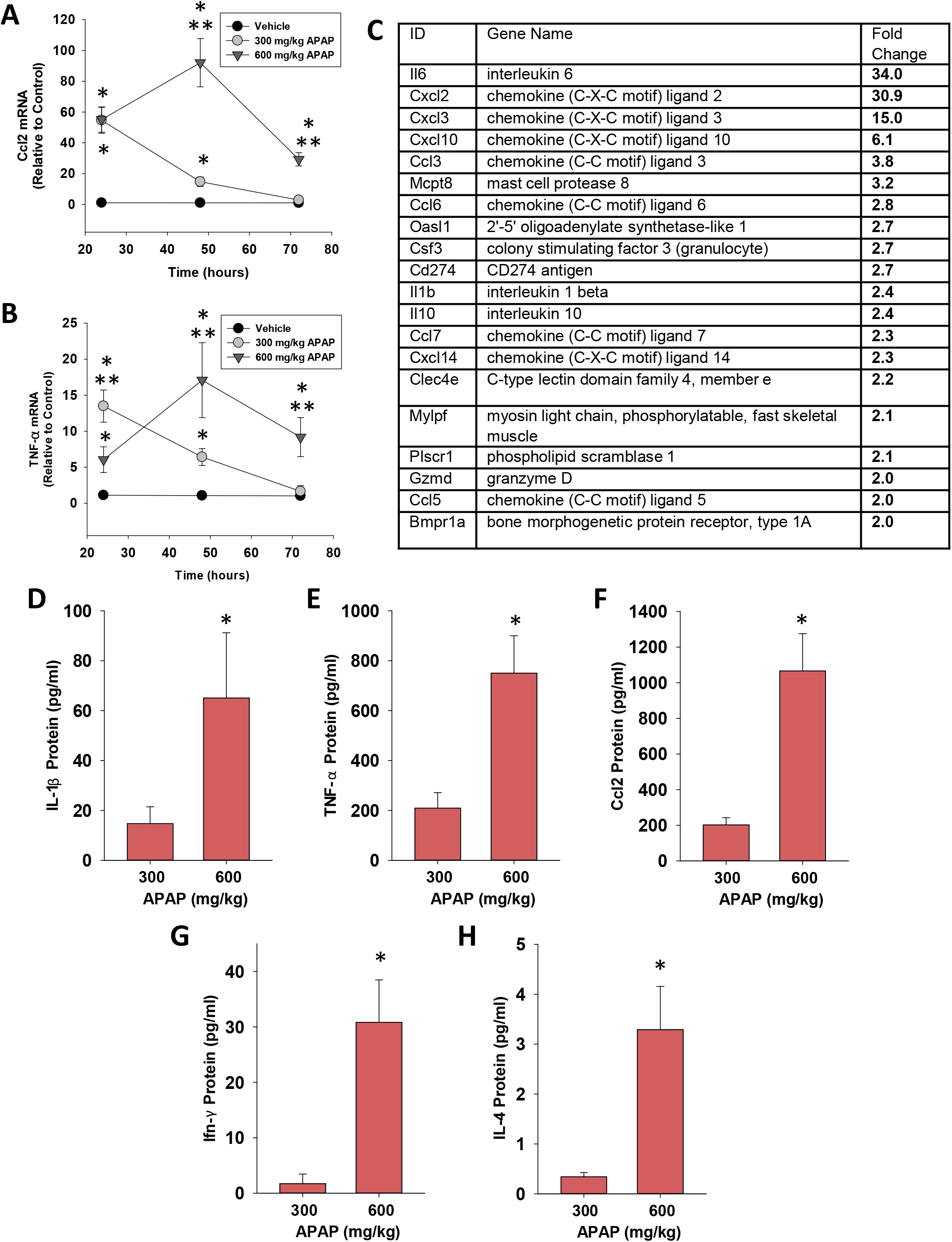
Quantification of cytokines in the livers and serum of mice treated with APAP. (A) Ccl2 and (B) TNF-α mRNA levels were measured at 24, 48, and 72 hours after APAP treatment. *Significantly different from vehicle-treated mice. **Significantly different from mice treated with 300 mg/kg at the same time point. Data are expressed as mean ± SEM; *n* = 5-10 mice per group. (C) mRNA levels of immunomodulatory genes increased in the livers of mice treated with 600 mg/kg APAP when compared to mice treated with 300 mg/kg APAP (*p*< 0.05). (D-H) Serum levels of the indicated cytokine were measured at 72 hours after APAP treatment. *Significantly different from mice treated with 300 mg/kg APAP. Data are expressed as mean ± SEM; *n* = 5-10 mice per group.

In ALF patients with SIRS that progresses to CARS, high systemic levels of proinflammatory cytokines frequently co-exist with high levels of anti-inflammatory cytokines, such as IL-10 ^14^. Similar to these clinical findings, hepatic mRNA levels of IL-10 were rapidly increased in mice treated with 600 mg/kg APAP and remained elevated at 72 hours (Fig. 4A). Importantly, elevated mRNA levels were matched by increased IL-10 protein in serum (Fig. 4B). By contrast, in mice treated with 300 mg/kg APAP, IL-10 mRNA levels did not increase until 72 hours after treatment (Fig. 4A and 4B). Analysis of purified populations of F4/80^+^ and Ly6C^+^ (i.e., MDMs) macrophages revealed elevated expression of IL-10 mRNA in F4/80^+^ cells but not Ly6C^+^ cells in mice treated with 600 mg/kg APAP (Fig 4C). Consistent with these findings, greater numbers of GFP-expressing F4/80^+^ cells were detected in the livers of IL-10 reporter mice treated with 600 mg/kg APAP (Fig. 4D and 4E). Collectively, these findings demonstrate the presence of both pro- and anti-inflammatory cytokines in mice treated with 600 mg/kg APAP similar to observations in ALF patients with SIRS and CARS.

**Figure 4.**
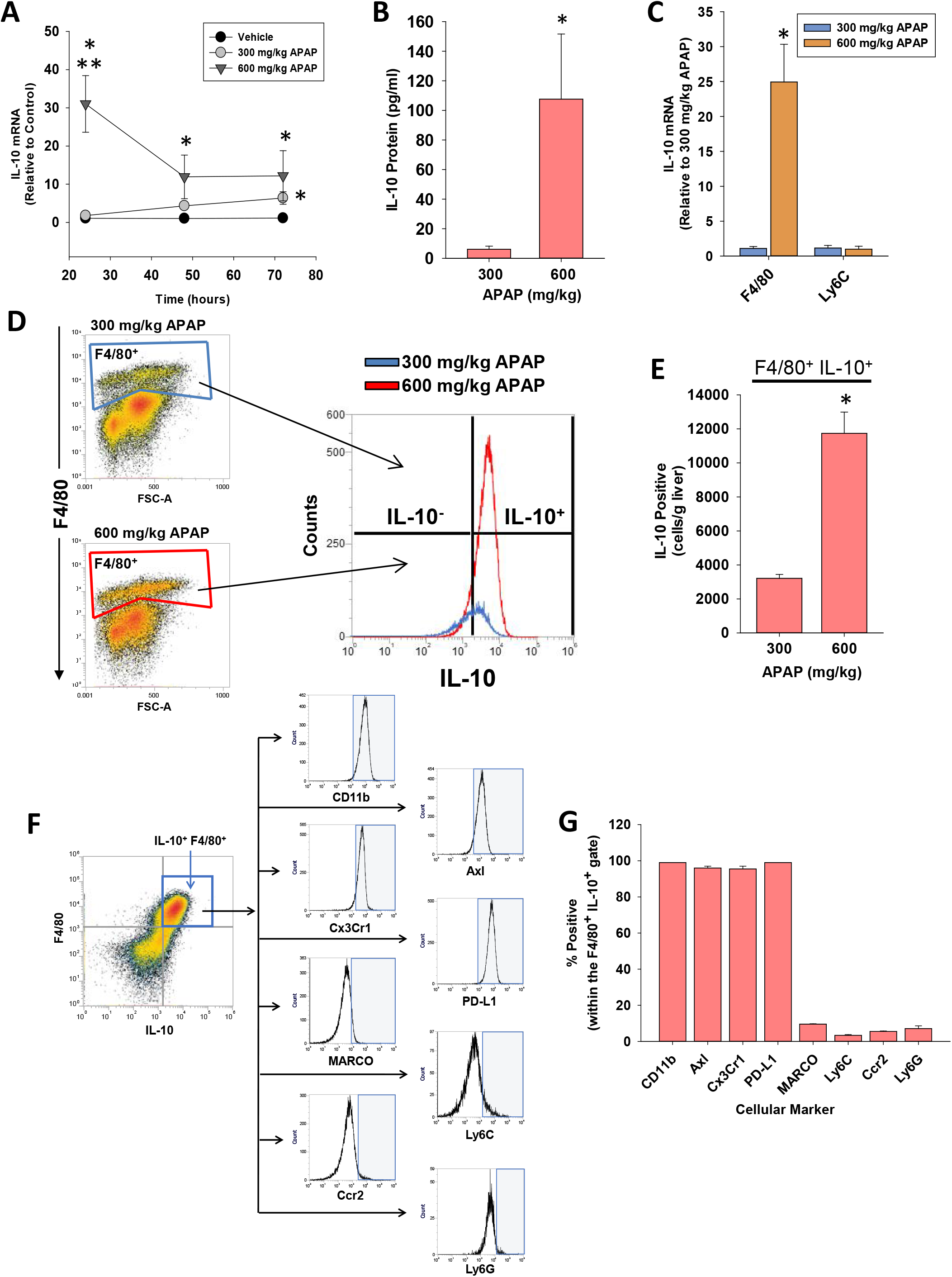
Hepatic and systemic levels of IL-10 in APAP-treated mice. Mice were treated with either 300 or 600 mg/kg APAP. (A) At the indicated time, IL-10 mRNA levels were measured in the liver. (B) IL-10 protein was measured in serum at 72 hours after APAP treatment. (C) F4/80^+^ and Ly6C^+^ myeloid cells were isolated from the liver at 24 hours after APAP, and IL-10 mRNA levels were measured. (D) Nonparenchymal cells were isolated from the livers of IL-10 reporter mice treated 24 hours earlier with 300 or 600 mg/kg APAP. Flow cytometry was used to identify IL-10 expressing F4/80^+^ cells. Gate for F4/80^+^ cells indicated in the density plots. Representative histogram of IL-10 expression in F4/80^+^ cells. (E) Quantification of the number of F4/80^+^ cells expressing IL-10 in the liver from flow cytometry. *Significantly different from mice treated with 300 mg/kg APAP. Data are expressed as mean ± SEM; n = 4-5 mice per group. (F) Mice were treated with 600 mg/kg APAP. At 24 hours later, flow cytometry was used to detect IL-10 expressing F4/80^+^ cells, indicated by the blue box. Representative histograms of IL-10^+^ F4/80^+^ cells expressing the indicated marker (x-axis). Positive staining indicated by the blue shaded box. (G) Quantification of the flow cytometry in F. Data are expressed as mean ± SEM; n = 4 mice per group.

### IMMUNOPHENOTYPING OF IL-10^+^ F4/80^+^ CELLS REVEALS A MYELOID-DERIVED SUPPRESSOR CELL-LIKE PHENOTYPE

IL-10 GFP reporter mice were treated with 600 mg/kg APAP for 24 hours. Flow cytometry was used to detect IL-10-expressing F4/80^+^ cells in the liver (Fig. 4F). Immunophenotyping of these cells revealed that they expressed CD11b, PD-L1, Cx3Cr1, and Axl consistent with a myeloid-derived suppressor cell (MDSC)-like phenotype (Fig. 4F and 4G). These cells did not express, however, additional markers commonly associated with MDSCs, including MARCO, Ly6C, Ccr2, or Ly6G (Fig. 4F and 4G).

### IL-10 PREVENTS TRAFFICKING OF MACROPHAGES INTO NECROTIC LESIONS IN MICE WITH APAP-INDUCED ALF

The mechanistic basis for a worse outcome in ALF patients with high systemic levels of IL-10 is not fully understood. Our earlier findings demonstrated that monocyte recruitment and trafficking were disrupted in the livers of mice treated with 600 mg/kg APAP, resulting in a failure to clear dead cell debris (Fig. 1 and 2). Because of the potent immune inhibitory properties of IL-10, we tested the hypothesis that IL-10 contributes to this defect. To examine this, we used both loss and gain of function approaches. In mice treated with 600 mg/kg APAP, injection of IL-10 neutralizing antibody, 24 hours after APAP, increased inflammatory cell infiltration into necrotic lesions by 72 hours (Fig. 5A-B and 5E-F). By contrast, in mice treated with 300 mg/kg APAP, pharmacological elevation of IL-10 24 hours after APAP, decreased inflammatory cell infiltration into necrotic lesions (Fig. 5C-D and 5H-I).

**Figure 5.**
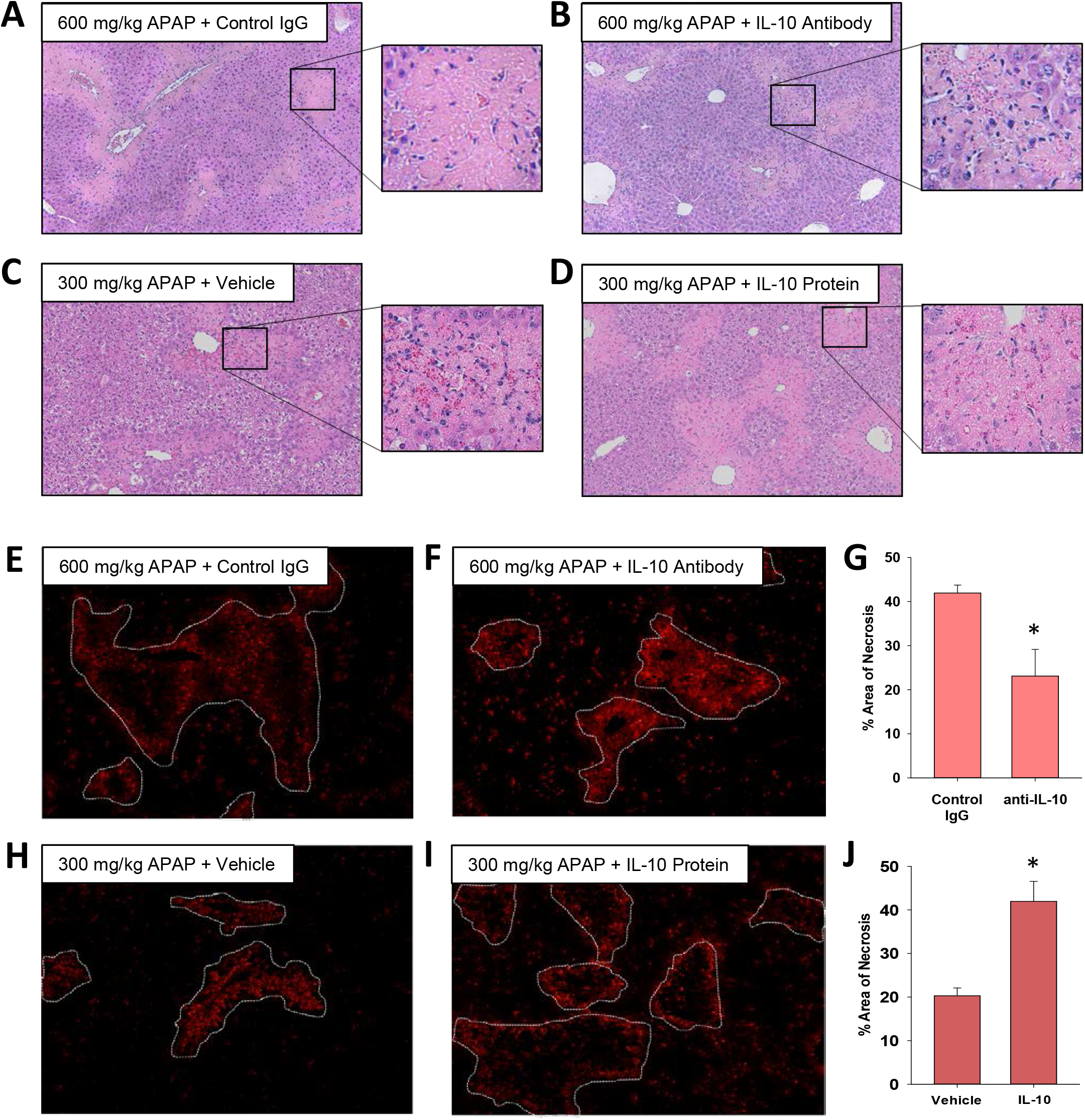
Impact of IL-10 on histopathology and inflammation in mice treated with APAP. (A-B and E-G) Mice were treated with 600 mg/kg APAP followed by treatment with control IgG or anti-IL-10 antibody 24 hours later. Livers were collected 72 hours after APAP treatment. (C-D and H-J) Mice were treated with 300 mg/kg APAP followed by treatment with vehicle or 5 mg recombinant IL-10 24 hours later. Livers were collected 48 hours after APAP treatment. (A-D) Representative photomicrographs of hematoxylin and eosin-stained liver sections. Box highlights a region of interest that is shown at a higher magnification. CD68 was detected by immunofluorescence in sections of liver from mice treated with (E) 600 mg/kg APAP and control IgG, (F) 600 mg/kg APAP and anti-IL-10 antibody, (H) 300 mg/kg APAP and vehicle or (I) 300 mg/kg APAP and recombinant IL-10. The necrotic lesions are demarcated by a dashed line. Representative photomicrographs from an n = 5 mice per group. (G and J) The area of necrosis was quantified. n = 5 mice per group. *Significantly different at p<0.05.

Importantly, restoration of MDM trafficking in mice treated with 600 mg/kg APAP and IL-10 neutralizing antibody was associated with a reduction in the area of necrosis by 72 hours (Fig 5G), whereas elevated levels of IL-10 in 300 mg/kg APAP mice increased the area of necrosis (Fig. 5J). Surprisingly, despite the effect on MDM recruitment and trafficking, modulation of IL-10 levels in mice treated with either 300 mg/kg APAP or 600 mg/kg APAP had minimal impact on the expression of proinflammatory cytokines (Supplemental Figs. 2 and 3).

Bhushan and colleagues demonstrated that hepatocyte proliferation is substantially lower in mice treated with 600 mg/kg APAP ^18^. To determine whether IL-10 is causally involved in this defect, we quantified PCNA positive hepatocytes. As shown in Supplemental Figures 2 and 3, modulation of IL-10 levels had no impact on hepatocyte PCNA staining.

### IL-6 CONTRIBUTES TO UPREGULATION OF IL-10 AND PD-L1 IN F4/80 EXPRESSING HEPATIC MACROPHAGES IN MICE WITH APAP-INDUCED ALF

Immunophenotyping of F4/80^+^ IL-10^+^ cells indicated an MDSC-like phenotype (Fig. 4). Studies have identified IL-6 as an important stimulus of MDSC formation in the tumor microenvironment, and IL-6 levels are highly elevated in patients with APAP-induced ALF ^10, 29^. Therefore, we next determined whether IL-6 levels are increased in mice treated with 600 mg/kg APAP and determined whether this is important for the generation of F4/80^+^ IL-10^+^ cells. Hepatic IL-6 mRNA levels were increased to a greater extent in mice treated with 600 mg/kg APAP when compared to mice treated with 300 mg/kg APAP at all time-points (Fig. 6A). The increase in IL-6 mRNA was matched by increased IL-6 protein in serum (Fig. 6B). IL-6 mRNA levels were greater in F4/80^+^ macrophages purified from the livers of mice treated with 600 mg/kg APAP, whereas IL-6 mRNA levels were not different in purified Ly6C^+^ cells (Fig. 6C). Next, mice were treated with IL-6 neutralizing antibody 2 hours after APAP injection to determine whether IL-6 contributed to the early induction of IL-10. As shown in Figure 6, neutralization of IL-6 reduced hepatic IL-10 mRNA levels by approximately 70% at 24 hours (Fig. 6D). Remarkably, neutralization of IL-6 reduced hepatic PD-L1 mRNA levels by greater than 90% (Fig. 6E). Previous studies demonstrated that liver injury is increased in IL-6 knockout mice treated with 300 mg/kg APAP ^30^. In mice treated with 600 mg/kg APAP, however, neutralization of IL-6 beginning at 2 hours after APAP treatment had no impact on liver injury (Fig. 6F). Unexpectedly, during these studies we observed a striking difference in the behavior of 600 mg/kg mice treated with IL-6 neutralizing antibody. Mice treated with 600 mg/kg APAP and control IgG experienced general paresis with significant gait abnormalities and demonstrated an apparent loss of balance when prompted to move. Surprisingly, 600 mg/kg APAP mice co-treated with IL-6 neutralizing antibody were more active and alert, were more responsive to stimuli, and did not display gait abnormalities when moving. Consistent with these behavioral observations, mortality was reduced in mice treated with IL-6 antibody (Fig. 6G).

**Figure 6.**
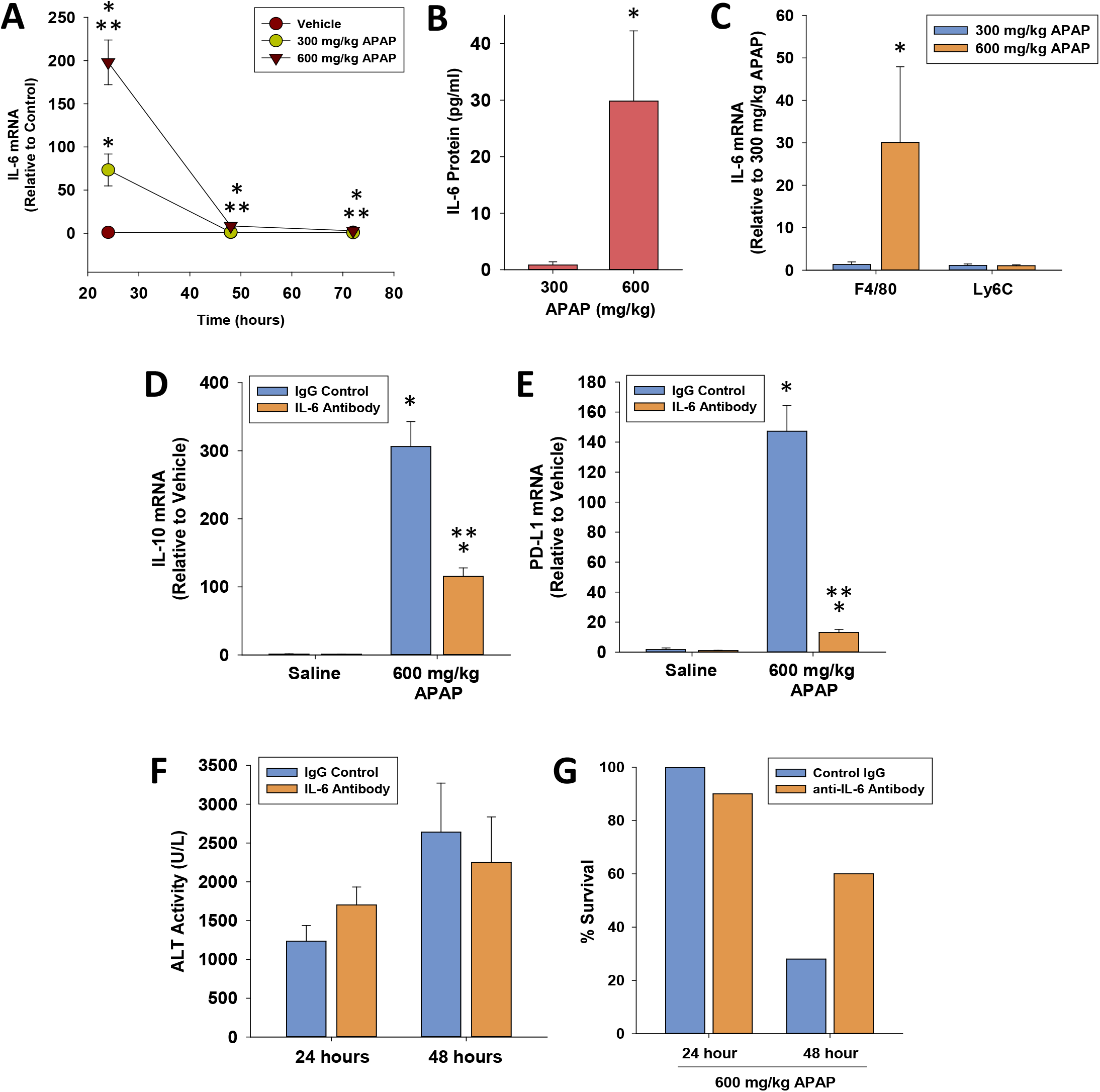
Role of IL-6 in regulation of IL-10 after APAP. Mice were treated with 600 mg/kg APAP at the indicated time. (A) IL-6 mRNA levels were measured at 24, 48, and 72 hours after APAP treatment. *Significantly different from vehicle-treated mice. **Significantly different from mice treated with 300 mg/kg at the same time point. (B) Serum levels of IL-6 were measured at 72 hours after APAP treatment. (C) IL-6 mRNA levels were measured in F4/80^+^ and Ly6C^+^ cells purified from the liver. *Significantly different from mice treated with 300 mg/kg APAP. Data are expressed as mean ± SEM; *n* = 5-10 mice per group. Mice were treated with 600 mg/kg APAP followed by IgG control or IL-6 neutralizing antibody 2 hours later. mRNA levels of (D) IL-10 and (E) PD-L1 were measured at 24 hours after APAP. (F) ALT activity at 24 and 48 hours. *Significantly different from vehicle-treated mice. **Significantly different from mice treated with 600 mg/kg and IgG control. Data are expressed as mean ± SEM; *n* = 3 mice per group for saline treated mice. *n* = 9 mice per group for mice treated with APAP and IgG control or IL-6 antibody. (G) Survival at the indicated time-point.

Rao and colleagues previously reported that mice treated with 500 mg/kg APAP display symptoms of hepatic encephalopathy, including lethargy and gait abnormalities, similar to what we observed in mice treated with 600 mg/kg APAP above ^31^. Further, they showed that ammonia, a proposed causative factor of hepatic encephalopathy, was increased in the serum and cerebral spinal fluid, and there was evidence of brain edema ^31^. Because IL-6 neutralization appeared to reduce symptoms of hepatic encephalopathy in our study, we examined whether IL-6 neutralization also impacted a more quantitative, less subjective endpoint. Prior studies reported that cerebral blood flow is reduced at early stages of hepatic encephalopathy in ALF patients ^32, 33^. Therefore, we examined whether cerebral blood flow is similarly reduced in mice with APAP-induced ALF and determined whether it was impacted by neutralization of IL-6. For this study, we used 500 mg/kg APAP since Rao and colleagues had previously used clinical assessments to determine the grade of hepatic encephalopathy in mice treated with this dose of APAP ^31^. Similar to mice treated with 600 mg/kg APAP, neutralization of IL-6 improved survival without affecting liver injury or hepatocyte proliferation in mice treated with 500 mg/kg APAP (Fig. 7A-7C). In mice treated with 300 mg/kg APAP, cerebral blood flow was not significantly impacted relative to vehicle controls (Fig. 7F and 7G). Consistent with findings of hepatic encephalopathy, however, total frontal blood flow was reduced in mice treated with 500 mg/kg APAP and control IgG (Fig. 7H). Remarkably, neutralization of IL-6 fully restored total frontal blood flow in mice treated with 500 mg/kg APAP, consistent with the marked differences in behavior in these mice (Fig. 7).

**Figure 7.**
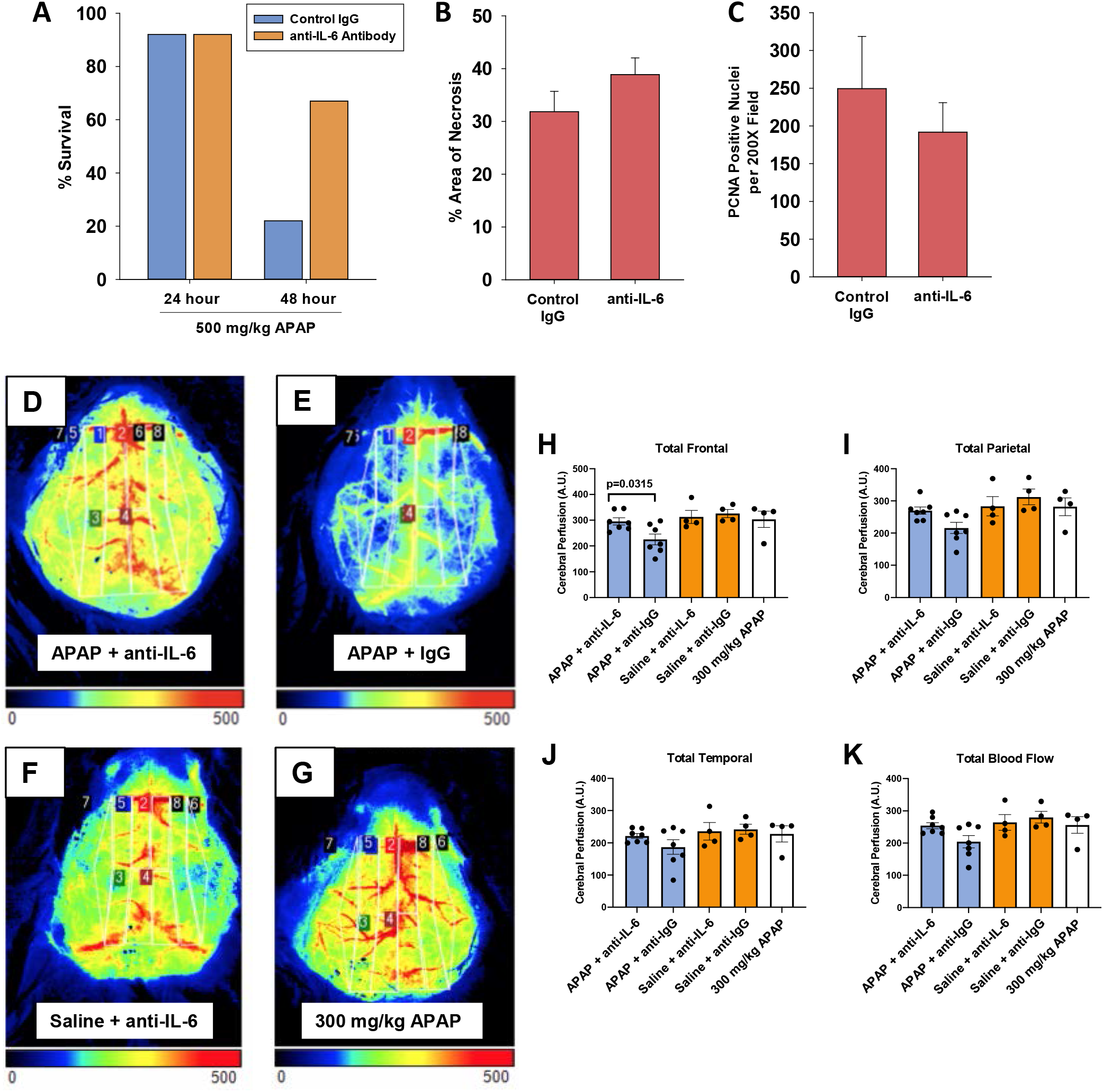
Role of IL-6 in liver injury and cerebral blood flow after APAP. Mice were treated with 500 mg/kg APAP followed by IgG control or IL-6 neutralizing antibody 2 hours later. (A) Survival at 24 and 48 hours after APAP. *n*=10 (B) Area of Necrosis and (C) PCNA positive cells were quantified at 24 hours after APAP. (D-K) Mice were treated with 500 mg/kg APAP followed by IgG control or IL-6 neutralizing antibody 2 hours later. Additional mice were treated with saline followed by IgG control or IL-6 neutralizing antibody 2 hours later. A separate cohort of mice were treated with 300 mg/kg APAP for comparison. Laser speckle imaging was used to quantify cerebral blood flow 24 hours after APAP. Representative brain images from laser speckle imaging from mice treated with (D) 500 mg/kg APAP + IL-6 neutralizing antibody, (E) 500 mg/kg APAP + Control IgG, (F) Saline + IL-6 neutralizing antibody, and (G) 300 mg/kg APAP. (H-K) Total cerebral blood flow and blood flow within the indicated brain region was quantified.

## Discussion

Several studies have reported a strong association between high levels of certain cytokines (e.g., IL-6 and IL-10) and the risk of multi-organ failure and death in ALF patients ^9, 10, 14^. While this has long been recognized in clinical settings, the mechanistic basis for cytokine dysregulation and its association with outcome in ALF remains poorly defined. Standard experimental settings of APAP overdose in mice (i.e., doses at or near 300 mg/kg) produce liver injury that is predictably repaired, triggering normalization of cytokine levels and resolution of inflammation. While studies using this approach have yielded important information regarding the mechanisms controlling cytokine synthesis and release during normal liver repair, they have not provided mechanistic insight into the pathogenesis of cytokine dysregulation occurring in ALF patients. Here we used a robust experimental setting of failed liver repair after APAP overdose in which a high dose of APAP was administered (i.e., 500-600 mg/kg to wild-type mice). This experimental approach, characterized in detail by Bhushan and colleagues, recapitulates many of the key features of ALF in critically ill patients, including impaired hepatocyte proliferation, renal injury, and evidence of hepatic encephalopathy ^18, 31, 34^. By using this approach, our studies reveal that cytokine dysregulation is a key feature of APAP-induced ALF in mice, recapitulating a critical aspect of ALF pathogenesis in patients. In support of this, high levels of several proinflammatory cytokines, resembling SIRS, were present in the serum of mice treated with 600 mg/kg at a time (i.e., 72 hours) where these cytokines were markedly lower in mice receiving 300 mg/kg APAP (Fig. 3). Further, high levels of IL-10 and PD-L1, a feature of CARS in ALF patients, also occurred coincident with high levels of proinflammatory cytokines, recapitulating a paradoxical feature of ALF in patients (Fig. 4) ^11, 14^. By using this pragmatic experimental approach, our studies identified specific cytokines that are causally involved in defective liver repair and in the development of certain extrahepatic complications that frequently arise in severe ALF. Furthermore, from these findings, we identified potential therapeutic targets that reduced mortality in mice and could conceivably reduce mortality in ALF patients with the poorest prognosis. Importantly, these targets of therapy would not have been apparent in mice treated with low doses of APAP. In fact, our studies indicate that beneficial, therapeutic interventions identified from low dose studies (e.g., stimulation of IL-6 signaling), may be detrimental to patients with severe ALF, further illustrating the importance of APAP dose for ALF studies.

Clinical studies have demonstrated a clear association between high systemic levels of IL-6 and severity of hepatic encephalopathy in ALF patients ^11, 12^. Our studies provide mechanistic support for this association by demonstrating that neutralization of IL-6 restores cerebral blood flow and reduces features of hepatic encephalopathy in mice with ALF (Fig. 7). Typically, cerebral perfusion is tightly regulated to address the high metabolic demands of the brain. Thus, the 25% reduction in cerebral blood flow observed in the ALF mice is significant. The hypoperfusion observed in the current study is similar to that seen in mice subjected to carotid artery occlusion or to prolonged periods of hypertension and cerebral small vessel disease indicating the true severity of this effect ^35, 36^. Although clinical studies have reported reductions in cerebral blood flow in ALF patients with hepatic encephalopathy, the importance of this to the pathogenesis of this disorder remains to be defined ^32, 33^. Our studies have identified an experimental setting that could be used to mechanistically interrogate this phenomenon in mice to examine whether there is a causal link between the reduction in cerebral blood flow and the acute cognitive and motor deficits that occur in hepatic encephalopathy.

Our studies indicate that Kupffer cells, the resident macrophages of the liver, may be the primary source of IL-6 and IL-10 in APAP-induced ALF (Fig. 4 and 6). Phenotypically, these cells resembled myeloid-derived suppressor cells (MDSCs) as indicated by high-level expression of IL-10, PD-L1, CD11b, Axl and Cx3Cr1 (Fig. 4) ^29^. MDSCs are highly immunosuppressive cells that have been extensively studied in cancer ^29^. While MDSCs have not been reported in patients with APAP-induced ALF, studies have demonstrated increased numbers of these cells in patients with acute-on-chronic liver failure ^37^. In these studies, higher numbers of MDSCs were associated with a worse outcome prompting the authors propose that the immune suppressive properties of these cells may produce a permissive environment for life-threatening infections to develop. Recently, Triantafyllou and colleagues reported that PD-1 and PD-L1 are increased on Kupffer cells in mice treated with APAP ^13^. Further, they demonstrated increased levels of PD-1 and PD-L1 on circulating monocytes in patients with APAP-induced ALF, suggesting that MDSCs may also be a feature of APAP-induced ALF ^13^. In this study, ALF patients with the highest levels of PD-1 and PD-L1 were at greatest risk of developing infections and had the highest mortality ^13^. In support of a causal role for PD-1 and PD-L1 in the increased susceptibility to infections, they demonstrated that anti-PD-1 treatment protected APAP-treated mice from severe sepsis induced by injection of live bacteria ^13^. Our study expands upon these findings and demonstrates further that IL-6 is critical for upregulation of PD-L1 in mice with APAP-induced ALF. Studies have shown that IL-6 can increase the immune suppressive activity of tumor-associated macrophages ^38^, suggesting that IL-6 may directly enhance the MDSC-like properties of Kupffer cells in APAP-induced ALF.

IL-10 is a potent anti-inflammatory cytokine that is hepatoprotective in several models of liver injury and disease. In fact, liver injury and mortality are greatly enhanced in IL-10 knockout mice treated with doses of APAP ranging from 120 to 300 mg/kg ^17^. This long-standing study has driven current dogma that the primary role of IL-10 in experimental liver damage is to inhibit inflammatory liver injury. Yet, these results do not recapitulate what is observed in ALF patients, where high levels of IL-10 are an independent predictor of a poor outcome ^14^. Because IL-10 is well known to suppress immunity against invading pathogens, it is possible that high levels of IL-10 in patients with ALF increases their susceptibility to infections that then progresses to sepsis. Interestingly, though, Berry and colleagues showed no association between high levels of IL-10 and incidence of sepsis in ALF patients ^14^, suggesting that IL-10 impacts additional processes that are critical for recovery from ALF. Our studies provide an alternative explanation for the connection between high IL-10 levels and poor outcome in ALF. Specifically, IL-10 negatively impacts the ability of MDMs to migrate within the liver. In mice treated with 300 mg/kg APAP, CD68^+^ macrophages filled the necrotic lesions by 48 hours (Fig. 2). By contrast, in mice challenged with 600 mg/kg APAP, recruited CD68^+^ macrophages were seemingly incapable of entering areas of necrosis, a phenomenon that has been noted in ALF patient livers (Fig. 2) ^11, 26^. Remarkably, in mice treated with 600 mg/kg APAP, neutralization of IL-10 completely restored macrophage trafficking into the necrotic foci (Fig. 5), and conversely, treatment of mice with recombinant IL-10, beginning at 24 hours after 300 mg/kg APAP, prevented intrahepatic macrophage trafficking into the necrotic lesions (Fig. 5). Collectively, these findings offer a mechanistic underpinning for the connection between high IL-10 levels and poor outcome in ALF.

The mechanistic basis for high IL-6 levels in mice treated with 600 mg/kg APAP is not fully clear, however, one potential stimulating factor may be severe hepatocellular hypoxia. Gao and colleagues recently showed that the hypoxia-regulated transcription factor, hypoxia-inducible factor-2α (HIF-2α), is activated in hepatic macrophages after treatment of mice with 300 mg/kg APAP ^30^. Further, they demonstrated that activation of HIF-2α in myeloid cells contributes to IL-6 upregulation ^30^. We and others have reported substantial sinusoidal congestion and hemorrhage in the livers of mice treated with 600 mg/kg APAP, a phenotype that is less severe in mice treated with 300 mg/kg APAP ^18^. It is possible that the severe sinusoidal congestion further impairs hepatic blood flow resulting in greater hypoxia, HIF-2α activation, and IL-6 upregulation.

Collectively, our findings demonstrate that myeloid cell dysfunction and cytokine dysregulation observed in ALF patients are effectively recapitulated in mice treated with a high dose of APAP. As such, this experimental setting provides a novel platform to interrogate mechanisms of immune dysregulation in ALF and to identify new therapeutic interventions. Further, by using this approach, our studies identified IL-6 as a central player in macrophage dysregulation in ALF, challenging the long-held belief that IL-6 is hepatoprotective in APAP-induced ALF and providing a mechanism to explain the paradoxical association between high levels of IL-6 and poor outcome in ALF patients. To our knowledge, no clinical studies have yet targeted IL-6 in ALF patients. Understandably, there may be reluctance to use this therapeutic approach considering the known hepatoprotective properties of IL-6 and the rare reports of acute liver injury occurring in patients receiving the anti-IL-6 antibody, tocilizumab ^39^. Further studies into the mechanism by which IL-6 becomes dysregulated in ALF, however, may identify an approach to “normalize” IL-6 levels thereby reducing the detrimental aspects of IL-6 signaling while maintaining its beneficial effects.

## Abbreviations

IL-10: interleukin-10
ALF: acute liver failure
APAP: acetaminophen
MDM: monocyte-derived macrophage
IL-1: interleukin-1
IL-6: interleukin-6
TNF-α: tumor necrosis factor-α
CCL2: C-C motif chemokine ligand 2
PCNA: proliferating cell nuclear antigen
PE: phycoerythrin
SEM: standard error of the mean
ANOVA: analysis of variance
IFN-γ: interferon-γ
IL-4: interleukin-4
MDSC: myeloid-derived suppressor cell
ACLF: acute on chronic liver failure
H&E: hematoxylin and eosin

---

**Supplemental Figure 1.**
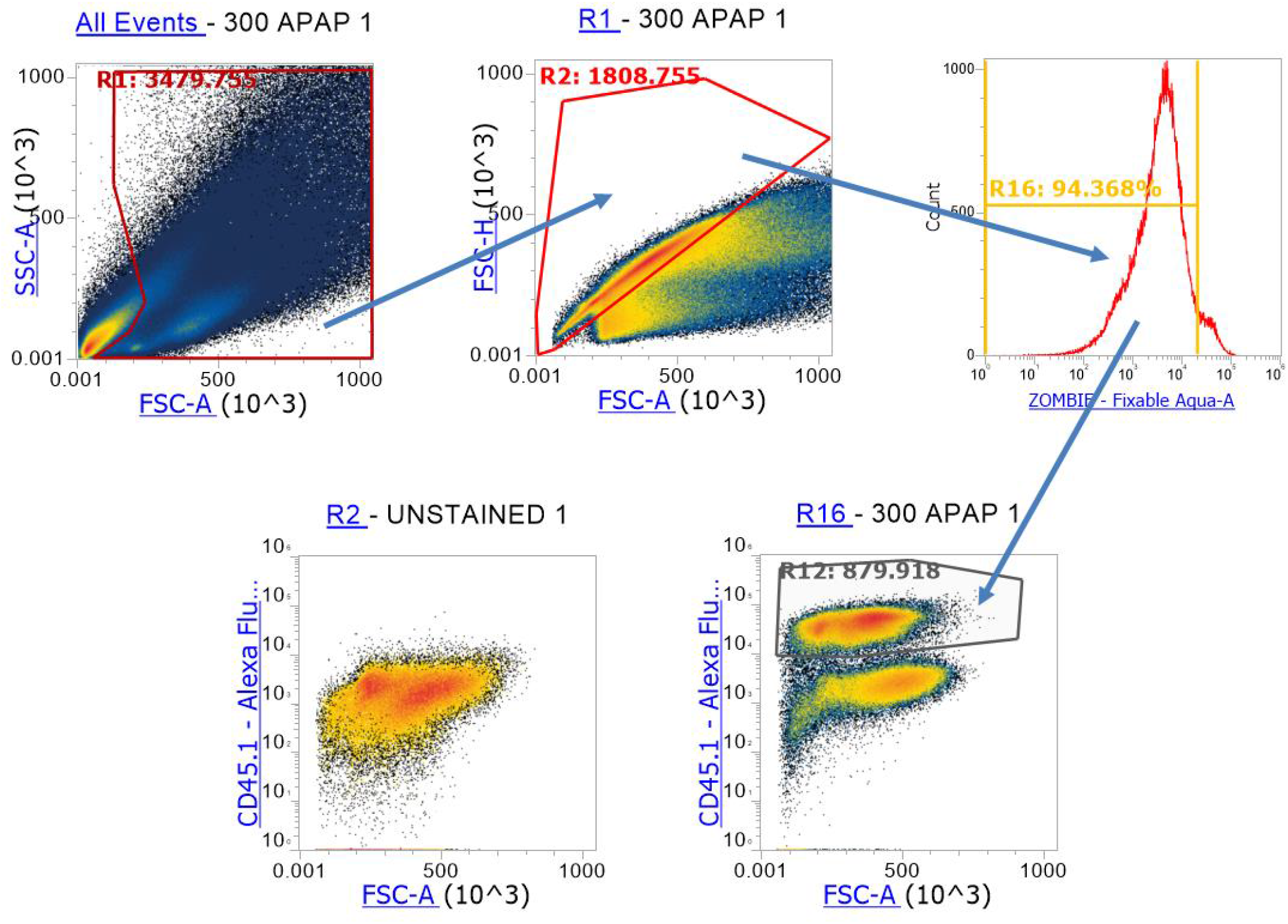
Gating strategy for flow cytometry.

**Supplemental Figure 2.**
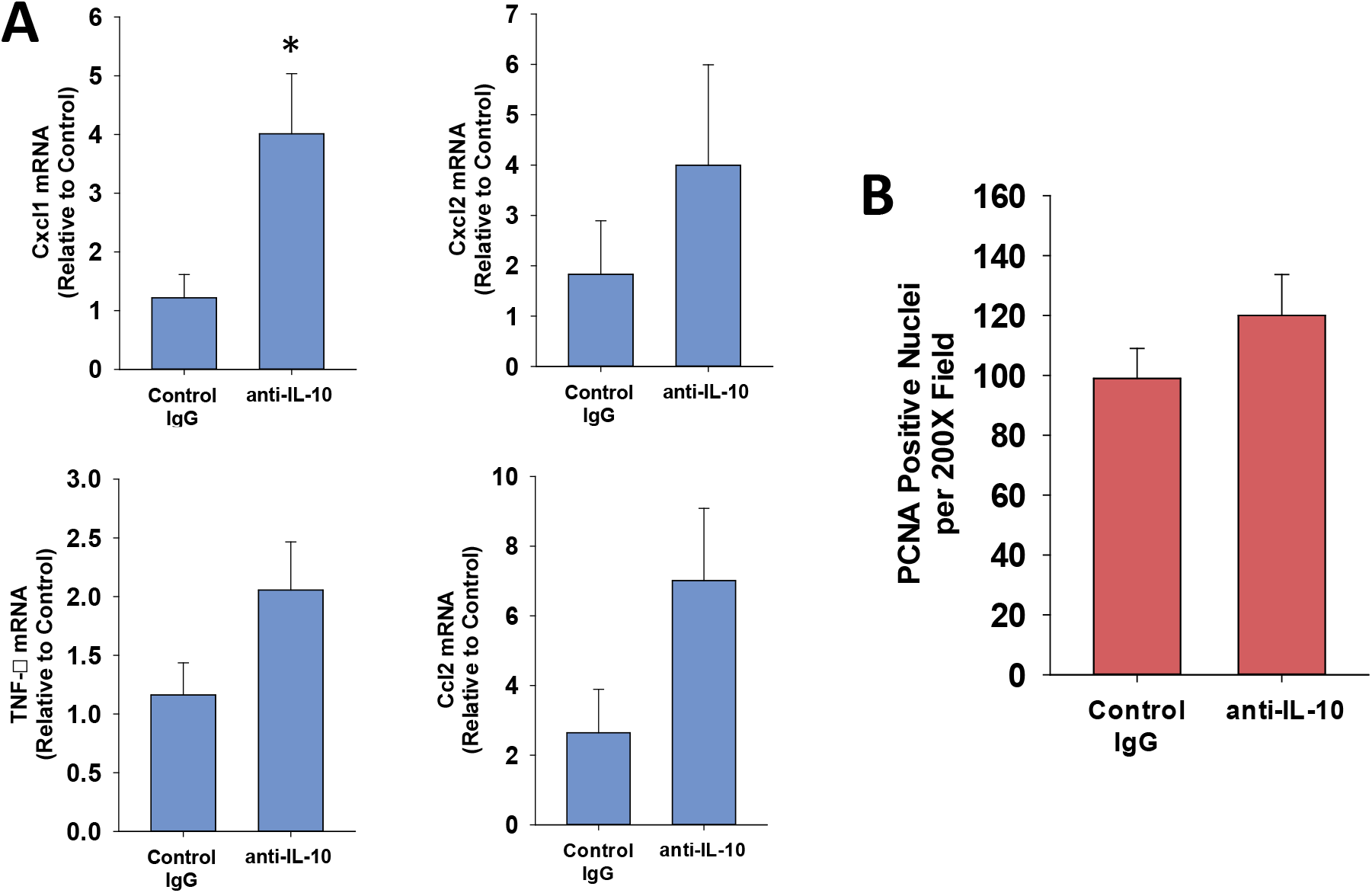
Impact of IL-10 on histopathology and inflammation in mice treated with APAP. (A-B) Mice were treated with 600 mg/kg APAP followed by treatment with control IgG or anti-IL-10 antibody 24 hours later. Livers were collected 72 hours after APAP treatment. (C-D) Mice were treated with 300 mg/kg APAP followed by treatment with vehicle or 5 mg recombinant IL-10 24 hours later. Livers were collected 48 hours after APAP treatment. (A and C) mRNA levels of selected cytokines in the liver. n=5. *Significantly different at p<0.05. (B and D) PCNA positive nuclei were quantified. n=5.

**Supplemental Figure 3.**
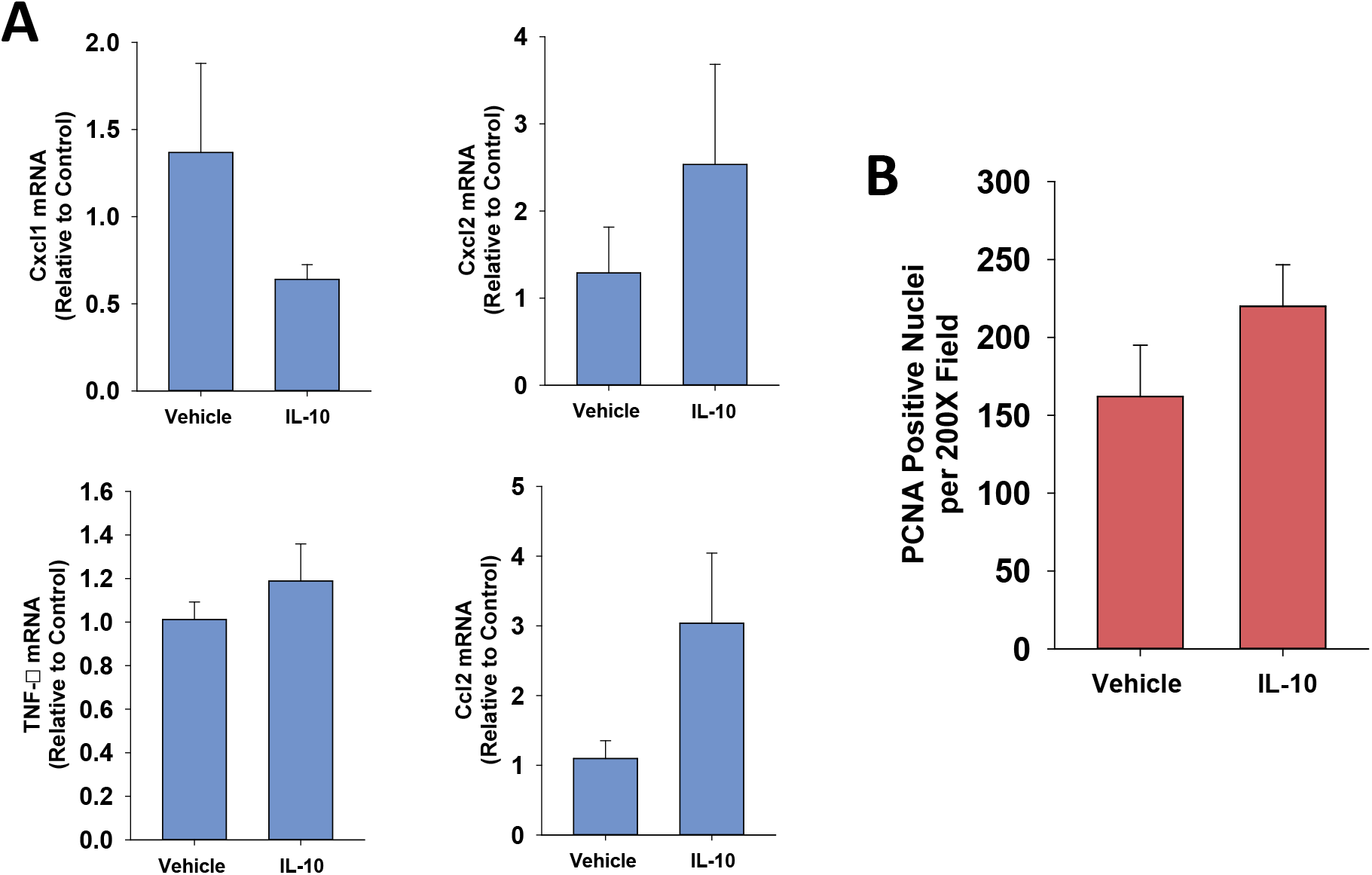
Impact of IL-10 on histopathology and inflammation in mice treated with APAP. (A-B) Mice were treated with 600 mg/kg APAP followed by treatment with control IgG or anti-IL-10 antibody 24 hours later. Livers were collected 72 hours after APAP treatment. (C-D) Mice were treated with 300 mg/kg APAP followed by treatment with vehicle or 5 mg recombinant IL-10 24 hours later. Livers were collected 48 hours after APAP treatment. (A and C) mRNA levels of selected cytokines in the liver. n=5. *Significantly different at p<0.05. (B and D) PCNA positive nuclei were quantified. n=5.

## References

1. Stravitz RT, Lee WM. Acute liver failure. Lancet 2019;394:869–881.

2. Lee WM, Squires RH, Nyberg SL, et al. Acute liver failure: Summary of a workshop. Hepatology 2008;47:1401–15.

3. Bernal W, Wendon J. Acute liver failure. N Engl J Med 2013;369:2525–34.

4. Saito C, Zwingmann C, Jaeschke H. Novel mechanisms of protection against acetaminophen hepatotoxicity in mice by glutathione and N-acetylcysteine. Hepatology 2010;51:246–54.

5. Smilkstein MJ, Knapp GL, Kulig KW, et al. Efficacy of oral N-acetylcysteine in the treatment of acetaminophen overdose. Analysis of the national multicenter study (1976 to 1985). N Engl J Med 1988;319:1557–62.

6. Prescott LF, Park J, Ballantyne A, et al. Treatment of paracetamol (acetaminophen) poisoning with N-acetylcysteine. Lancet 1977;2:432–4.

7. Lee WM. Recent developments in acute liver failure. Best Pract Res Clin Gastroenterol 2012;26:3–16.

8. Stravitz RT. Critical management decisions in patients with acute liver failure. Chest 2008;134:1092–1102.

9. Rolando N, Wade J, Davalos M, et al. The systemic inflammatory response syndrome in acute liver failure. Hepatology 2000;32:734–9.

10. Antoniades CG, Berry PA, Davies ET, et al. Reduced monocyte HLA-DR expression: a novel biomarker of disease severity and outcome in acetaminophen-induced acute liver failure. Hepatology 2006;44:34–43.

11. Antoniades CG, Berry PA, Wendon JA, et al. The importance of immune dysfunction in determining outcome in acute liver failure. J Hepatol 2008;49:845–61.

12. Butterworth RF. Pathogenesis of hepatic encephalopathy and brain edema in acute liver failure. J Clin Exp Hepatol 2015;5:S96–S103.

13. Triantafyllou E, Gudd CL, Mawhin MA, et al. PD-1 blockade improves Kupffer cell bacterial clearance in acute liver injury. J Clin Invest 2021;131.

14. Berry PA, Antoniades CG, Hussain MJ, et al. Admission levels and early changes in serum interleukin-10 are predictive of poor outcome in acute liver failure and decompensated cirrhosis. Liver Int 2010;30:733–40.

15. James LP, Lamps LW, McCullough S, et al. Interleukin 6 and hepatocyte regeneration in acetaminophen toxicity in the mouse. Biochem Biophys Res Commun 2003;309:857–63.

16. James LP, McCullough SS, Lamps LW, et al. Effect of N-acetylcysteine on acetaminophen toxicity in mice: relationship to reactive nitrogen and cytokine formation. Toxicol Sci 2003;75:458–67.

17. Bourdi M, Masubuchi Y, Reilly TP, et al. Protection against acetaminophen-induced liver injury and lethality by interleukin 10: role of inducible nitric oxide synthase. Hepatology 2002;35:289–98.

18. Bhushan B, Walesky C, Manley M, et al. Pro-regenerative signaling after acetaminophen-induced acute liver injury in mice identified using a novel incremental dose model. Am J Pathol 2014;184:3013–25.

19. Gregory B, Larson AM, Reisch J, et al. Acetaminophen dose does not predict outcome in acetaminophen-induced acute liver failure. J Investig Med 2010;58:707–10.

20. Mochizuki A, Pace A, Rockwell CE, et al. Hepatic stellate cells orchestrate clearance of necrotic cells in a hypoxia-inducible factor-1α-dependent manner by modulating macrophage phenotype in mice. J Immunol 2014;192:3847–3857.

21. Allen K, Jaeschke H, Copple BL. Bile acids induce inflammatory genes in hepatocytes: a novel mechanism of inflammation during obstructive cholestasis. Am J Pathol 2011;178:175–86.

22. Kim ND, Moon JO, Slitt AL, et al. Early growth response factor-1 is critical for cholestatic liver injury. Toxicol Sci 2006;90:586–95.

23. Bhushan B, Gunewardena S, Edwards G, et al. Comparison of liver regeneration after partial hepatectomy and acetaminophen-induced acute liver failure: A global picture based on transcriptome analysis. Food Chem Toxicol 2020;139:111186.

24. Holt MP, Cheng L, Ju C. Identification and characterization of infiltrating macrophages in acetaminophen-induced liver injury. J Leukoc Biol 2008;84:1410–21.

25. Zigmond E, Samia-Grinberg S, Pasmanik-Chor M, et al. Infiltrating monocyte-derived macrophages and resident kupffer cells display different ontogeny and functions in acute liver injury. J Immunol 2014;193:344–53.

26. Antoniades CG, Quaglia A, Taams LS, et al. Source and characterization of hepatic macrophages in acetaminophen-induced acute liver failure in humans. Hepatology 2012;56:735–46.

27. Mossanen JC, Krenkel O, Ergen C, et al. Chemokine (C-C motif) receptor 2-positive monocytes aggravate the early phase of acetaminophen-induced acute liver injury. Hepatology 2016;64:1667–1682.

28. Roth K, Strickland J, Joshi N, et al. Dichotomous Role of Plasmin in Regulation of Macrophage Function after Acetaminophen Overdose. Am J Pathol 2019;189:1986–2001.

29. Yaseen MM, Abuharfeil NM, Darmani H, et al. Mechanisms of immune suppression by myeloid-derived suppressor cells: the role of interleukin-10 as a key immunoregulatory cytokine. Open Biol 2020;10:200111.

30. Gao RY, Wang M, Liu Q, et al. Hypoxia-Inducible Factor-2α Reprograms Liver Macrophages to Protect Against Acute Liver Injury Through the Production of Interleukin-6. Hepatology 2020;71:2105–2117.

31. Rama Rao KV, Verkman AS, Curtis KM, et al. Aquaporin-4 deletion in mice reduces encephalopathy and brain edema in experimental acute liver failure. Neurobiol Dis 2014;63:222–8.

32. Bjerring PN, Gluud LL, Larsen FS. Cerebral Blood Flow and Metabolism in Hepatic Encephalopathy-A Meta-Analysis. J Clin Exp Hepatol 2018;8:286–293.

33. Wendon JA, Harrison PM, Keays R, et al. Cerebral blood flow and metabolism in fulminant liver failure. Hepatology 1994;19:1407–13.

34. Akakpo JY, Ramachandran A, Orhan H, et al. 4-methylpyrazole protects against acetaminophen-induced acute kidney injury. Toxicol Appl Pharmacol 2020;409:115317.

35. Salinero AE, Robison LS, Gannon OJ, et al. Sex-specific effects of high-fat diet on cognitive impairment in a mouse model of VCID. FASEB J 2020;34:15108–15122.

36. Diaz-Otero JM, Fisher C, Downs K, et al. Endothelial Mineralocorticoid Receptor Mediates Parenchymal Arteriole and Posterior Cerebral Artery Remodeling During Angiotensin II-Induced Hypertension. Hypertension 2017;70:1113–1121.

37. Zeng Y, Li Y, Xu Z, et al. Myeloid-derived suppressor cells expansion is closely associated with disease severity and progression in HBV-related acute-on-chronic liver failure. J Med Virol 2019;91:1510–1518.

38. Jiang M, Chen J, Zhang W, et al. Interleukin-6. Front Immunol 2017;8:1840.

39. Mahamid M, Paz K, Reuven M, et al. Hepatotoxicity due to tocilizumab and anakinra in rheumatoid arthritis: two case reports. Int J Gen Med 2011;4:657–60.

